# Origins of *cin*: Lateral Gene Transfer of Cytoplasmic Incompatibility Nuclease Operon to *Orientia tsutsugamushi*

**DOI:** 10.1101/2025.03.04.641471

**Authors:** Kyle Oswalt, Seun Oladipupo, Luis Mendez, Joseph J. Gillespie, John F. Beckmann

## Abstract

CinB nucleases are *Wolbachia* proteins that induce cytoplasmic incompatibility (CI) through tandem nuclease domains nuc_1_ and nuc_2_ ^1^. CI is a form of reproductive parasitism (RP) whereby males are conditionally sterilized ^2^. The system behaves as a toxin-antidote (TA) system ^1–8^ where operon gene *cinA* encodes an antidote and gene *cinB* encodes a toxin ^1,5^*. Cin* operons are purportedly the cause of gene-drive induced by wolbachiae infecting *Drosophila simulans* ^1,9–12^. An unanswered research question is whether lateral transfer of CI operons to bacteria outside wolbachiae would transfer RP phenotype and activate gene-drive. We demonstrate that a *cin* operon, capable of gene-drive, has jumped into the genome of *Orientia tsutsugamushi*, a human pathogen and causative agent of lethal scrub typhus. When expressed in transgenic *Drosophila melanogaster*, the wildtype *cin*B^*o*Tsu^ was capable of inducing CI independent of its partner antidote *cin*A^*o*Tsu^. In addition, *cin*A^*o*Tsu^ rescued the phenotype in accordance with strict TA functionality. To understand the diverging roles of the tandem nuclease domains we mutated the domains and tested all permutations of active/inactive forms. Finally, we isolated *IS*5 transposon variants flanking the operon in *O*. *tsutsugamushi* and re-activated them to test their mobility. We demonstrate that these transposons can transfer genes and initiate lateral gene transfers into *E. coli*. These data demonstrate that active bacterial transposons can mobilize and transfer CI factors (*cifs*) to diverse bacteria. Overall, our data contribute mechanistic understanding in support of the TA model of CI and illuminate biochemical mechanisms that mobilize *cifs* across genomes from phylogenetically diverse taxa.

**Significance Statement:** CI operons are foundational genes that directly contribute to the success of *Wolbachia*-based bio-control strategies. Two applications of RP-inducing *Wolbachia* strains are insect population replacement and the incompatible insect technique. Both these techniques do not work if *cifs* do not function. Thus, a mechanistic understanding of *cif* function contributes to worldwide bio-control implementations. Furthermore, certain wolbachiae have long been studied as broad-spectrum inducers of diverse RP phenotypes, including CI, parthenogenesis, male-killing, and feminization. How these diverse phenotypes evolve and if they are all induced by *cif* genes is a long-standing field question. For example, *Orientia tsutsugamushi*, the causative agent of a deadly scrub typhus, induces parthenogenesis in *Leptotrombidium* mites, which putatively have reproductive advantages over uninfected mites. How *Orientia* induces RP is an important epidemiological question in vector biology. In our report, we show that an active *cin* operon is capable of CI and jumped into *Orientia* genomes via an *IS*5 transposon. We reconstructed this transposon and engineered it as a biotechnological tool. In toto, our study leads to the proposal of a new hypothesis whereby CI and parthenogenesis phenotypes might both be connected to the same *cif* expressed under divergent host genetic contexts.

## Introduction

Reproductive parasitism (RP) comes in many forms, including cytoplasmic incompatibility (CI) and parthenogenesis ^13^. In both cases, RP skews Mendelian inheritance to spread the RP inducer. Thus, the outcome of RP is the same as gene-drive and, as such, we view it as a form of natural gene-drive. In the literature, RP “inducers” are sourced to microbial endosymbionts (i.e., some species/strains of *Wolbachia, Spiroplasma,* and *Cardinium*) that live inside arthropod host gonads ^14–16^. However, a more basal view of RP focuses on the actual genes that induce it rather than the organism. Many characterized RP inducer genes are demonstrably laterally transferred ^17–23^. Therefore, an RP phenotype might spread from one host to another via lateral gene transfer (LGT).

A major RP phenotype, CI, is controlled by toxin-antidote (TA) operons ^1,3–8,24–26^. These genes are broadly termed CI factors (*cifs*), or *cid*, *cin*, *cnd*, depending upon the molecular mechanism of CI induction ^1,3,4,25,27,28^. Various LGT mechanisms have been proposed to mobilize *cifs*. The operons putatively shuffle amongst *Wolbachia* endosymbionts (Alphaproteobacteria: Rickettsiales: Anaplasmataceae), via their WO-phages, transposons and other mobile elements; though precise molecular jumping mechanisms haven’t been demonstrated in the laboratory ^5,20,21,23,25,27,29^. Transposons, as mobile genetic elements, are known to facilitate LGT. Thus, transposons may contribute to the spread of RP phenotypes to different bacterial taxa. This is especially concerning in cases where transposons transfer genetic cargo into pathogens. The potential for transposon-mediated gene transfer could allow these RP phenotypes to expand into new contexts.

The identification of *Wolbachia* CI operons ^24,25^ led to their bioinformatic discovery outside wolbachiae. Some species of *Rickettsia*, which are in the Rickettsiales family (Rickettsiaceae), carry a *cnd* operon on a mobile plasmid, including the human pathogen *R*. *felis*^21^. Additionally *cin* operons are present in strains of another deadly Rickettsiaceae pathogen, *Orientia tsutsugamushi* ^21,30^, the etiological agent of lethal Scrub Typhus ^30–32^. Other intracellular bacteria, many with unknown human pathogenicity and/or unidentified RP phenotypes, also harbor *cin* and *cnd* operons ^21,33^. In toto, *cif* distribution patterns demonstrate frequent lateral gene transfer across diverse intracellular microbes (LGT)^18,20,21^. Because *cins* induce RP,^1^ we hypothesize that *cin* transfer might also transfer RP phenotypes to intracellular pathogens, particularly those that infect arthropod gonads.

*O. tsutsugamushi* is a neglected tropical disease ^30–32^ that annually infects over a million people ^34,35^. Much of the *biology of Rickettsiales* pathogens is similar to *Wolbachia*. For instance, like wolbachiae*, O. tsutsugamushi* infects ovaries of arthropods. *O. tsutsugamushi* has perfected maternal transmission and is vertically passed at rates near 100% ^36^. The vectors of *O. tsutsugamushi* are the larval (“chigger”) stages of mites in the genus *Leptotrombidium* (family Trombiculidae). Trombiculids are related to velvet mites, like the spider mite *Tetranychus urticae*, with ∼87% amino acid identity ^37,38^. Curiously, a *Wolbachia* strain that induces CI exists *in T. urticae* ^39–42^ but isn’t reported in Trombiculids. In contrast, Trombiculid mites infected with *O. tsutsugamushi* exhibit facultative thelytokous parthenogenesis, where females hatch from unfertilized eggs ^40–42^. When infected with *O. tsutsugamushi,* two vectors of scrub typhus, *L. fletcheri* and *L. arenicola*, produced only females, whereas uninfected colonies produced normal female:male ratios of ∼2:1 ^41,42^. In 1977, researchers hypothesized that the inducer of RP within these infected Trombiculids could be a common factor shared amongst *O. tsutsugamushi*, wolbachiae, and *Spiroplasma* species ^41,43,44^. Now that the genetic factors of RP have been determined in the latter two organisms, the hypothesis remains feasible given the prodigious exchange of distantly related RP inducing deubiquitylases (DUBS) and nucleases amongst arthropod endosymbionts ^21,28,45^. Transposons may also contribute to the mobility of these RP-inducing elements, facilitating the spread of RP phenotypes across species. A disconcerting thought is *Leptotrombidium* mites acquiring RP phenotypes via LGT of *cifs* to their *O. tsutsugamushi* populations; whereby they might displace non-infected mites and expand endemic regions of scrub typhus ^40^. This could lead to more human deaths. Our paper points out evidence where this may have happened in evolutionary history.

Another open question is whether *cifs* might be the source of other forms of RP beyond CI. *Wolbachia* are capable of parthenogenesis (in various mechanistic forms), feminization, and male-killing ^13^. A single strain can switch RP phenotype upon transfection to novel hosts ^46^. It’s logical that various RP phenotypes might be linked to the same inducer gene and we sought to begin exploring this. Parthenogenesis is the most complex RP phenotype. The term is, in some ways, a misnomer used to describe no less than four distinct molecular mechanisms including thelytoky, arrhenotoky, deuterotoky, and gynogenesis ^47^. Thelytoky, the exclusive production of females from unfertilized eggs, is the form of RP induced by *O. tsutsugamushi*. Thelytoky can itself be subdivided into approximately four distinct mechanisms that co-occur in *Acari* (see **Discussion**). Thus, the term parthenogenesis encompasses many discrete bio mechanisms and is likely not specific enough as a scientific term to be useful. Whether or not some of these bio mechanisms source to *cifs* in mites and/or other arthropods remains to be determined. With caution, *cin* function might be sufficient to explain a few mechanistic cases. To complicate matters, the specific type of parthenogenesis in *Leptotrombidium* vectors is unknown, but due to the seriousness of pathogen gene-drive via RP mechanisms, we investigated the function of this strange *cin* operon that jumped to *O. tsutsugamushi*, hereafter *cin*^oTsu^.

## Results

### *Orientia tsutsugamushi* Acquired Original *cin*^oTsu^ via Lateral Gene Transfer

Previously, we determined that the *O*. *tsutsugamushi* str. Boryong genome encoded a probable *cin* operon ^21^. Here, we demonstrate that highly similar *cin*-like operons are encoded in eight recently published *O*. *tsutsugamuishi* genomes ^48^ as well as the previously sequenced *O*. *tsutsugamushi* str. Ikeda genome. Remarkably, nearly all *cin* operons reside in regions of degraded Rickettsiales Amplified Genetic Elements (RAGEs), which are unique integrative conjugative elements carried by certain *Rickettsia* species and all sequenced strains of *O*. *tsutsugamushi* ^49–52^ (**Figure 1a**). *Rickettsia* RAGEs insert in tRNA genes and are usually single copy in genomes (or sometimes multicopy or mostly degraded) ^52–54^. In contrast, *Orientia* RAGEs are fragmented, with chunks of varying size numerously copied throughout the genome, making scrub typhus genomes extraordinarily repetitive ^49,51^. Despite subtle differences in “cargo genes” that piggyback on RAGE elements, *Rickettsia* and *Orientia* RAGEs are more similar to one another than to any other observed conjugative elements ^52^. Aside from certain species of *Tisiphia* (a recently named genus that is sister to genus *Rickettsia* ^55,56^), no other Rickettsiales genomes have been shown to carry RAGEs. Curiously, certain cargo genes on *Rickettsia* RAGEs have high homology to genes encoded in *Wolbachia* prophage genomes ^52^, leading to our earlier hypothesis that RAGEs and/or their transposon cargo facilitate lateral gene transfer (LGT) of CI genes across intracellular microbes ^21^. In summary, *cin* operons are present within genetic regions of high mobility.

**Figure 1.**
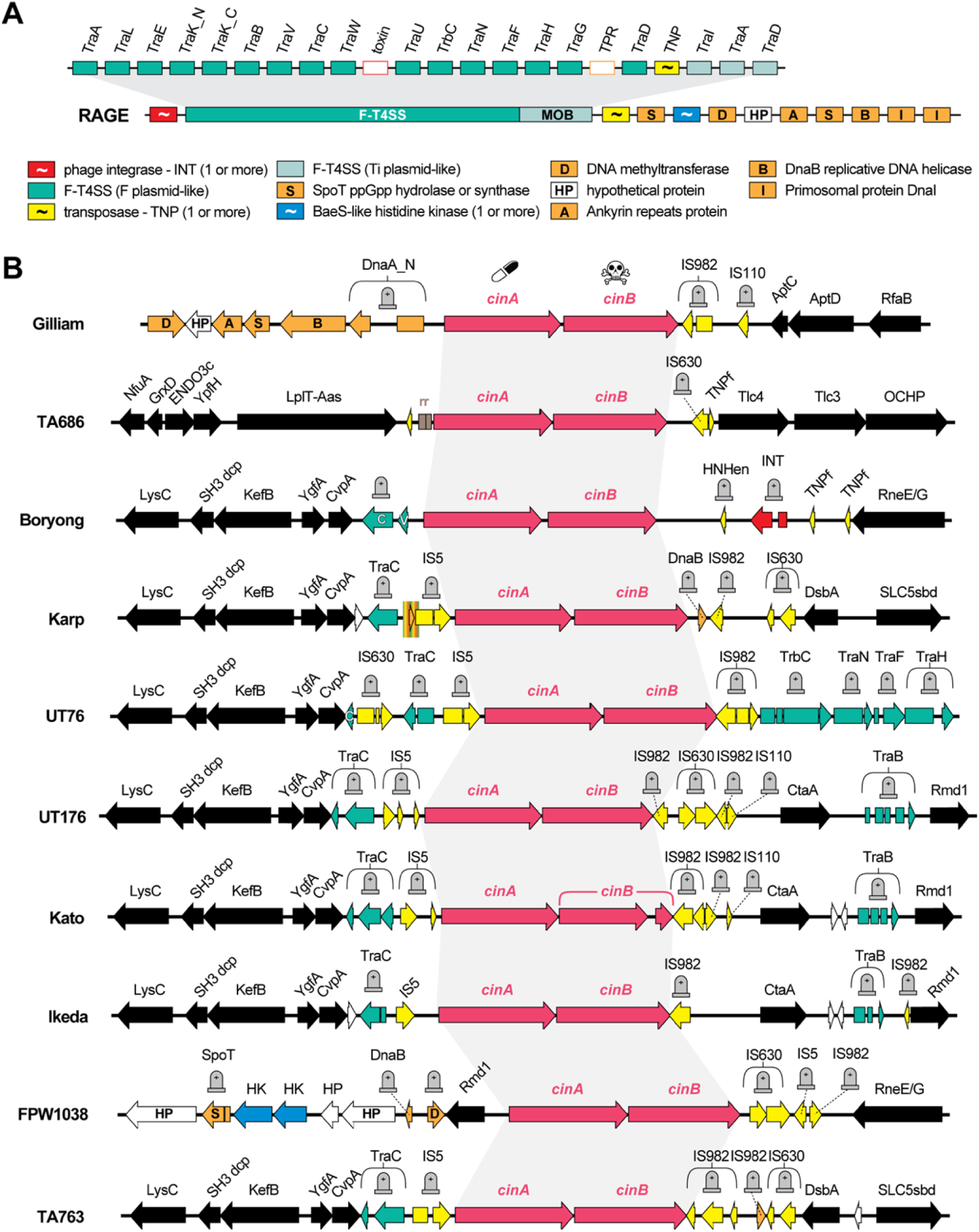
*Orientia tsutsugamushi* genomes encode *cin* operons associated with mobile genetic elements. **a.** A generalized model for the Rickettsiales Amplified Genetic Elements (RAGEs) present in *O*. *tsutsugamushi* genomes ^71–74^. In contrast to complete RAGEs in certain *Rickettsia* genomes, *O*. *tsutsugamushi* genomes possess massive proliferation of RAGE fragments interspersed with conserved genes. **b.** The predicted *cin* operon previously detected in the *O*. *tsutsugamushi* str. Boryong genome ^75^ was found to occur on chromosomes of nine sequenced *O*. *tsutsugamushi* genomes. *cin* operons are flanked by RAGE genes or other mobile genetic elements (see panel **a.** for color scheme). Conserved genes are black. For the *O*. *tsutsugamushi* str. Karp genome, orange markers depict a partial duplication of the DnaB gene indicating transposon-mediated insertion within this locus. Grave stones indicate broken pseudogenes. Pink genes indicate the *cin* operon.

All *Orientia cin* operons have definable insertion points near RAGE loci (**Figure 1b**). The *cin* operons are present only once in each genome, with flanking conserved genes indicating a probable single inheritance and minor rearrangement mediated by proliferated RAGE fragments. The genomes indicate that a *cin* operon either inserted 1) directly into a resident *O*. *tsutsugamushi* RAGE or 2) inserted into a primitive RAGE that then integrated into an ancestral *O*. *tsutsugamushi* genome. The *O*. *tsutsugamushi* str. Karp genome evidences a transposon-mediated insertion of the *cin* operon that duplicated part of a DnaB gene fragment (**Figure 1b**, rainbow swatch). Most importantly, the *IS*5 element possesses two inverted repeats (IRs) flanking the transposon, as well as an IR downstream of the neighboring *cin* operon. The presence of these specific IRs is strong bioinformatic evidence that the *IS*5 transposon carried the *cin* operon as cargo into *Orientia* genomes.

We ^5721,58^ and other researchers ^29^ previously posited that smaller transposons likely moved CI operons across diverse obligate intracellular bacteria. This hypothesis contrasts with phage derived movement ^59^, which seems unlikely in this scenario because larger mobile elements like RAGE and *Wolbachia* phage are genus restricted in contrast to these particular transposons ^52^. In addition, *Orientia* genomes lack complete phages, yet harbor viral-like genes similar to *Wolbachia* gene transfer agents ^60,61^. Investigative phylogenomics analyses of CinA and CinB protein sequences identify the most similar counterparts in *Rickettsia* genomes, namely *R*. *hoogstraalii* and *R*. *amblyommmatis* (**Sup. Fig.1a**). The *Rickettsia* CI-like operons generally contain larger toxins with multiple domains (e.g., CidB-like deubiquitinase domains, ankryrin repeats, latrotoxin domains, etc.) outside of the nuclease domains (**Sup. Fig. 1b**). Thus, this *cin* variant likely moved from rickettsiae into *O*. *tsutsugamushi*, given that we previously showed the smaller and simplified *cif* toxins likely evolved from the gene fission of larger, multi-domain *cif* toxins^62^.

### *cin*^oTsu^ Has a Unique Polymorphism in the Nuclease Catalytic Triad

Given the importance of *cin* to CI, we further inspected the *cinB* gene. Specifically, *cinB* alleles possess two PD-(D/E)XK nuclease domains (nuc_1_ and nuc_2_, see **Fig. 2**). These domains share a predicted secondary structure, αβββαβ, within a charged DEK catalytic triad ^1,5,25,63^. Chen et al. specifically characterized CinB*^w^*^Pip^ as a DNA nuclease. The triad catalyzes cleavage of DNA phosphate backbones by coordinating two Mg^2+^ ions and two water molecules toward nucleophilic attack ^1^. Chen et al. convincingly demonstrated that both nuc_1_ and nuc_2_ domains are required in CinB*^w^*^Pip^ to generate toxicity in yeast models. Furthermore, disabling nuc_2_ in transgenic CI constructs weakened CI ^1^. One unusual feature of native *cin*^*o*Tsu^ is that with respect to domains nuc_1_ and nuc_2_, which are conserved in other CinB toxins, it exists in a state of active-On/inactive-Off (On/Off) respectively (**Figure 2a**). The lysine to alanine mutation in the DEK catalytic triad is expected to ablate function of that nuc_2_ domain.

**Figure 2.**
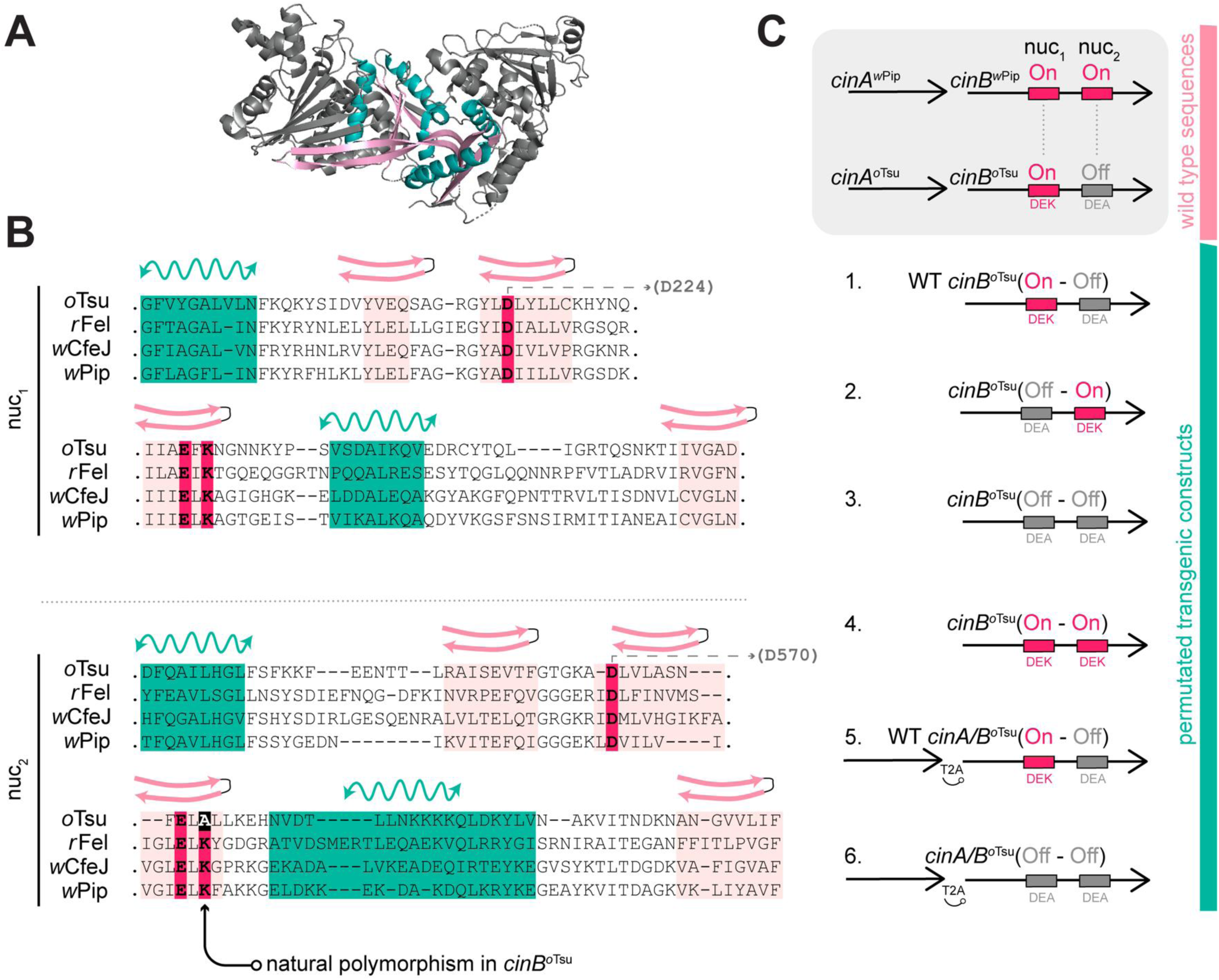
Comparison and design of *cin*B^*o*Tsu^ genetic constructs. **a.** 3-D structure of cinB*^w^*^Pip 76^**b.** Alignment of CinB^*o*Tsu^ with characterized CinB proteins. Green α-helices and pink β-sheets mark αβββαβ secondary predictions. Domains nuc_1_ and nuc_2_ are the N-terminal and C-terminal nuclease domains, respectively. Hot pink bars show conserved DEK catalytic triads. An arrow points to the natural DEA catalytic polymorphism in CinB^*o*Tsu^. **c.** Design of permutated nuclease constructs. Grey box shows native *cin* operons from *w*Pip and *Orientia tsutsugamushi* (*o*Tsu). Pink nucleases are “On”, meaning conserved relative to other characterized nuclease domains, and grey nucleases are “Off”, meaning divergent from other characterized nuclease domains. T2A is a viral peptide that induces ribosomal skipping and translation of two proteins from a single open reading frame ^45^.

Because *cin*^*o*Tsu^ is an evolutionary edge case outside the characterized wolbachiae toxins, as well as most other CinB proteins from other microbes, we felt this polymorphism might inform on the evolution of CI within *O. tsutsugamushi* and *Wolbachia*. We also questioned whether the *cin*^*o*Tsu^ was actively capable of RP in *Leptotrombidium* mites, though this has not been tested. Therefore, we undertook the task of testing all different combinatorial permutations of nuc_1_ and nuc_2_ in On or Off status. Recognizing that transgenic *D. melanogaster* is a robust model of *cif* function,^1,5,6,27,64–66^ we designed six different constructs to test transgenic CI phenotypes (**Figure 2b**). We cloned the six permutated *cin*^oTsu^ constructs into the integration vector pUASp-attb ^67^ and used ΦC31 mediated integration to insert the transgenes onto chromosome 3 of *D. melanogaster*.

### *cin*^oTsu^ Induces Cytoplasmic Incompatibility in Transgenic *Drosophila melanogaster*

To test for transgenic CI, we made the various UAS-transgene flies homozygous and crossed them into the maternal triple driver (MTD) strain to induce expression and measured hatch rates of transgene bearing males and females^66^ (**Figure 3**). When we expressed wild type (WT) *cinB*^*o*Tsu^(On – Off) we saw complete CI. With the reciprocal mutated construct, *cinB*^*o*Tsu^(Off – On), we saw minor CI (**Figure 3a**). These results are surprising for a few reasons. First, prior models suggested that the *cin*A partner would be required to induce CI according to the 2 x 1 model (i.e., that CI induction in males requires co-expression of two factors—a toxin and its partner—while rescue in females requires only the antidote), ^64,68^ but our data suggest this model does not apply to *cinB*^*o*Tsu^. Secondly, our data suggest that one intact nuclease domain (either nuc_1_ or nuc_2_) is sufficient for CI induction in this unique *cif* nuclease toxin. When both nuclease domains are inactivated CI was not observed.

**Figure 3.**
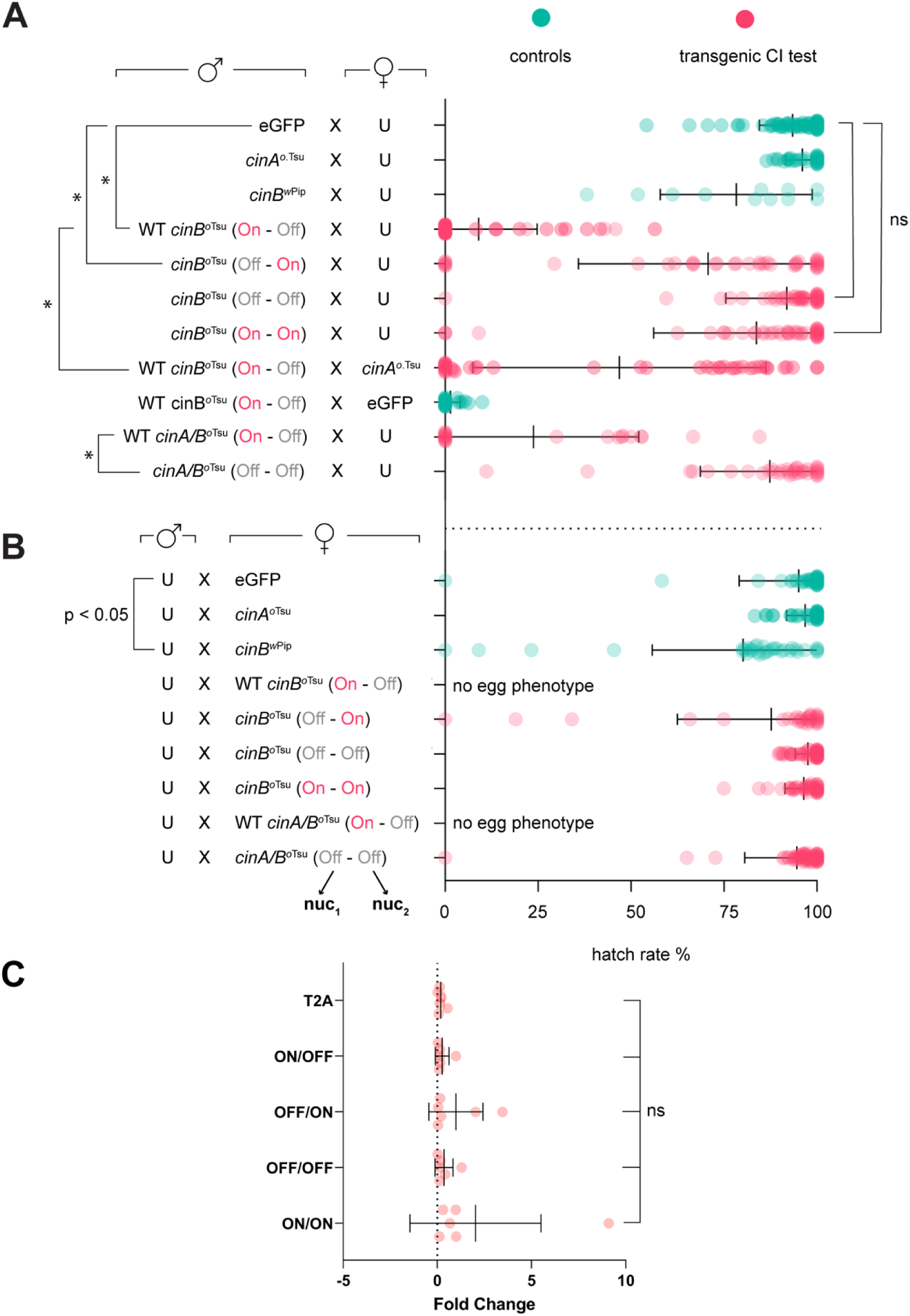
Hatch-rate analysis of transgenic *cin*^*o*Tsu^ constructs. Green data are control experiments and pink data are novel transgenic construct tests. **a.** Crosses with transgenes in males mated to uninfected mothers (U); or mothers with *cin*A^*o*Tsu^ or eGFP in rescue experiments. Lines are means with standard deviation. **b.** Crosses of uninfected males mated to females expressing transgenes. Significance (*) is p < 0.0001 and ns is not significant by ANOVA with multiple comparison between all groups and Tukey’s post-hoc analysis. GAL4/UAS driver is MTD. **c.** Relative gene expression of CinBott mutations in *Drosophila* fly lines using RT-qPCR. Fold change in gene expression of the OTT_0690 *O*. *tsutsugamushi* CinB-like gene was measured in five fly lines: T2A, ON/OFF, OFF/ON, OFF/OFF, and ON/ON. Expression levels were normalized to a housekeeping gene (Actin 5C). Bars represent the mean ± SEM from biological replicates. Statistical significance between fly lines was assessed by Kruskal-Wallis test with Dunn’s post-hoc test. Significance is indicated by p < 0.05.

Curiously when we tested the mutated *cinB*^*o*Tsu^(On – On) configuration we did not see CI. This suggests that the function of *cinB*^*o*Tsu^ likely differs from that of *cin*B*^w^*^Pip^ and that the wild type nuclease configuration of On – Off nuclease domains has some functional purpose. These differences might inform on how different RP phenotypes evolve in diverse arthropod hosts of reproductive parasites. Importantly, when we crossed WT *cinB*^*o*Tsu^(On – Off) to transgenic mothers overexpressing *cinA*^*o*Tsu^, we observed a strong bimodal rescue effect. The operon behaves differently than others tested and for the first time we observed the bimodal rescue pattern. About half the crosses were completely rescued and the other half did not rescue at all (a mean around 50% hatch rate).

We also tested expression of the full WT *cinA/B*^*o*Tsu^(On – Off) operon together with its partner gene and again saw CI with weaker effect, though this reduction by the presence of *cinA*^*o*Tsu^ was not statistically significant. Similarly, inactivation of the nuc_1_ and nuc_2_ domains in the full operon ablated this phenotype as a negative control. Because *cinA/B*^*o*Tsu^ behaved unlike other tested *cifs*, we also tested all reciprocal crosses by placing transgenes in females (**Figure 3b**). When we expressed WT *cinB*^*o*Tsu^(On – Off) we noticed that all females produced zero eggs. All other non-wildtype permutations did not show the “no eggs” phenotype. We again saw the no eggs phenotype when expressing the native operon, and again it disappeared when we inactivated both nuc_1_ and nuc_2_ domains within the operon. RT-PCR analysis confirmed that transgenic phenotypes could not be attributed to differential levels of transgene expression (**Figure 3c**).

### Transgenic Permutations of *cin*B^oTsu^ Induce Sex Specific Phenotypes

Because we observed a no egg phenotype and had never seen this before amongst prior *cif* studies, and because fecundity effects are not directly measured in hatch-rate graphs, we additionally began measuring fecundity (**Sup. Fig. 2)** ^1,5,6^. When expressing transgenes in females, non-native *cinB*^*o*Tsu^ transgenes did not differ from control mothers expressing a control, eGFP. Only WT *cinB*^*o*Tsu^(On – Off) showed major and significant reductions in fecundity, whether constituted independently or with its partner antidote within the operon (**Sup. Fig. 2a**). Our control *cin*B*^w^*^Pip^ only showed a weak reduction in fecundity that was significant, suggesting that fecundity effects are different amongst variable *cin*B toxins. These data suggest that the native sequence of *cinB*^*o*Tsu^(On - Off) is a general toxin with no sex specificity. This is a novel observation in *cifs* where most CinB toxins induce male specific sterility.

We hypothesize that this unique *cif* analog exhibits some intermediate or unique RP phenotype that has not yet effectively evolved sex specific sterilization of more mature CI systems; or perhaps it induces forms of parthenogenesis that current transgenic systems cannot assess in mites. We also tested whether transgenic males crossed into uninfected wild type females showed any fecundity effects (**Sup. Fig. 2b**). These data are harder to interpret, yet a key observation was that when the full WT *cin*A/B^oTsu^ (On – Off) operon was expressed in males and mated to uninfected wild type, it conferred severe and significant reductions in fecundity to that female; this implies that the transgenic proteins were transferred to females through mating and impacted female fecundity. This effect was reversed when the operon was catalytically inactivated in the cinA/B^oTsu^ (Off - Off) construct and was not even observed when just the toxin alone was expressed in males. We posit that CinA^oTsu^ helps assist CinB^oTsu^ during escape from sperm in females. Such a packaging role for CinA has been suggested prior (but before escaping sperm) ^4,6^ and this unique *cif* could prove as a useful contrast in future sperm/egg localization studies^69^.

### Maternal Transgenic Expression of *cinB*^oTsu^ Destroys Ovaries

The WT *cinA/B*^oTsu^ (On - Off) operon is clearly a powerful nuclease effector with broadly toxic effects in both males (CI) and females (reduction in fecundity); this is expected of general toxins. To further understand reductions in fecundity we used microscopy to visualize ovaries expressing WT *cinA/B*^oTsu^ (On - Off) (**Sup. Fig.3)**. All ovaries expressing this transgene (60/60) showed grossly stunted ovaries, deformed oviducts, or non-symmetrical ovarioles when compared to normal uninfected flies. We conclude that overexpression of WT *cinA/B*^oTsu^ (On - Off) in transgenic females reduces fecundity by shear destruction of their ovaries. These data replicate a phenotype heretofore only observed in transgenic *Aedes* mosquitos ^8^.

### Design and Reconstruction of the *Orientia IS*5 Transposon Constructs

Despite rampant pseudogenization and recombination, the *IS*5 transposon was determined to immediately flank the *cinA/B* operon in most *O*. *tsutsugamushi* genomes (**Fig 1b**); in addition, IRs of this mobile element indicated lateral gene transfer of the operon. Therefore, we sought to verify the potential of this *IS*5 transposon’s capability to laterally transfer genes. Bioinformatic analysis revealed that the *IS*5 transposon was pseudogenized due to three premature stop codons, rendering the transposase inactive. To resurrect its functionality, we strategically corrected these mutations to restore the open reading frame (ORF), resulting in a repaired sequence referred to as WT*. To assess the transposon’s ability to mobilize genetic material, we designed constructs featuring both the resurrected WT* transposase and a synthetic, codon-optimized version intended to enhance expression in *E. coli*, our heterologous host (**Sup. Fig 4**). Another unknown that needed to be determined was the complete length of the IR sequences. Each transposase variant was combined with two different lengths of IRs flanking the *kanR* antibiotic resistance cassette, which served as a cargo gene to evaluate transposition efficiency.

The first test set of IRs, termed “Long IRs,” were the native wild-type sequences from in the *O. tsutsugamushi* strain Karp (117 base pairs, bp). The second test set, called “Short IRs,” were synthetically designed sequences (17 bp), informed by the suggestion that *IS5* transposons prefer IRs of length 10-30 base pairs ^70^. By testing both long and short IR lengths and comparing the resurrected WT* transposase with the synthetic, codon-optimized version, we systematically investigated the key elements necessary for transposon activity. The genetic system relies on a mobilizable *kanR* cassette flanked by IRs, which allowed us to monitor successful transposition events through kanamycin resistance; this assay system quantifies transposition efficiency as colony-forming units (CFUs) and enabled us to assess the impact of both IR length and transposon sequence on transposition activity in *E. coli* (**Fig 4a-b**.).

**Figure 4.**
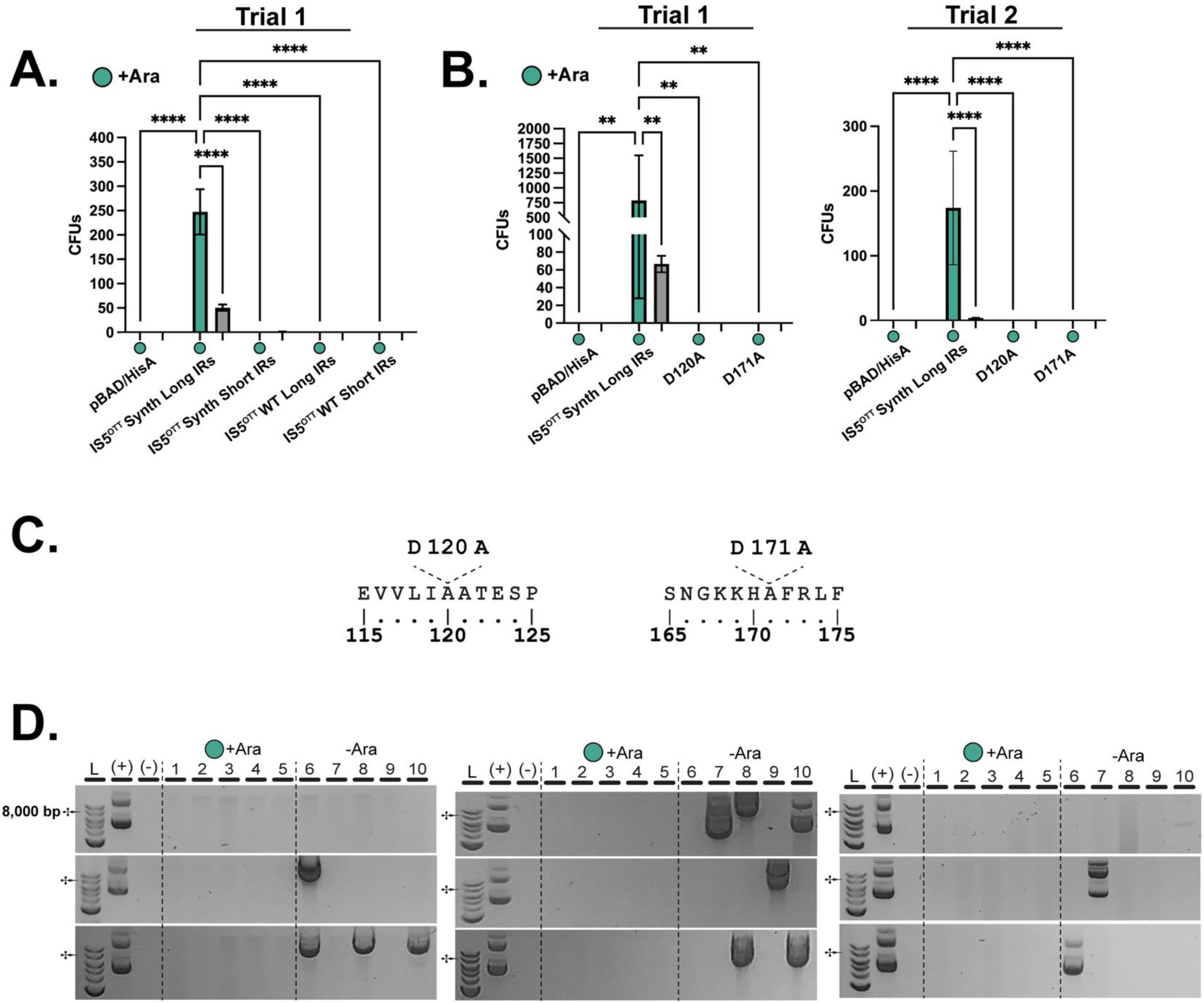
Evaluation of transposon activity and plasmid loss in *E. coli* BL21-AI. **a** CFU counts showing transposon activity across different strains under arabinose-induced conditions. The *IS*5^OTT^ Synth Long IRs strain shows the highest CFU counts. Statistical significance is indicated by asterisks (****p < 0.0001, ***p < 0.001, **p < 0.01). **b** CFU counts showing transposon activity of *IS*5^OTT^ Synth Long IRs and catalytic mutants. Catalytic mutants show significant decreases in activity. **c** Mutations of the catalytic sites of *IS*5^OTT^ Synth. **d** Agarose gel electrophoresis of minipreps from selected clones post-counter-selection. The 7,131 bp plasmid is indicated by the 8,000 bp marker. Lanes 1–5 (+Ara) show results under arabinose-induced conditions, while lanes 6–10 (−Ara) represent non-induced conditions. Positive (+) and negative (−) controls are included. DNA ladders are shown on the left.

We observed a significant increase in CFUs exclusively with the *IS*5^OTT^ Synth Long IRs construct upon arabinose induction (**Fig. 4a**). Our findings demonstrated that the synthetic (codon-optimized) transposase combined with the WT Long IRs functioned, mobilizing the *kanR* cassette under selective conditions (**Fig 4a**). In contrast, constructs with Short IRs or the wild-type *Orientia* DNA sequence exhibited markedly reduced or negligible transposition activity in *E.coli*. These results highlight the importance of both transposase sequence optimization and IR length in engineering efficient bacterial transposition. Following initial trials we investigated suspected catalytic sites belonging to a suspected DDE Endonuclease Domain (**Sup. Fig 5b**) found within the WT sequence of *IS*5^Ott^. As a negative control, two catalytic mutations (D120A and D171A) were induced via site directed mutagenesis and assessed (**Fig. 4c**). Catalytic inactivating mutants showed transposon inactivation when compared to the active controls (**Fig 4b**).

### Mapping and Characterizing Inserts

The elevated CFU counts correspond to colonies that underwent both transposition and plasmid loss during counterselection with 2-deoxy-D-galactose (2-DOG). Confirming plasmid loss was crucial to ensure that kanamycin resistance resulted from genomic integration rather than plasmid retention, which could lead to false positives. Selected colonies were subjected to plasmid minipreps, verifying the absence of the plasmid (**Fig. 4d**). Plasmid loss data shows that negative control colonies that grow on counter selective media have the chance of being a false positive due to plasmid retention as evident by banding present in negative lanes (**Fig. 4d**). As trials were conducted in recA (+) BL21AI, perhaps this background false positive rate could be attributed to recombination events inactivating galK on the plasmids.

Following counterselection and CFU screening, we sequenced the genomes of selected colonies with Illumina whole genome sequencing to confirm the integration of the Kanamycin resistance (*kanR*) cassette into the BL21-AI genome. The sequencing results verified successful transposon-mediated insertion, and that the Long IRs were maintained during the process. This led to the accurate transfer of a 1.1 kb segment into multiple loci within the host genome. A map shows the nine independent clonal insertions where the *kanR* cassette inserted into genes including *dtpD*, *gspE*, *speD*, and *malS* amongst others **(Fig. 4**). We evaluated all gene knockouts in *E. coli* using the EcoCyc database and determined that all genes are non-essential in *E. coli* per previous mutational analyses.

Further analysis of the flanking sequences at these integration sites revealed the expected sequence duplications were correctly generated during transposition **(Sup.6 Fig. 5a**). The functional analysis of the synthetic *IS*5^OTT^ Synth transposase (**Sup. Fig 5**) includes a sequence logo depicting the nucleotide composition around the insertion sites and a schematic of the transposition event. The sequence logo highlights the transposase’s preference to insert two positions downstream of an adenine (A) and directly upstream of a thymine (T). During this process, the base N immediately downstream of the A is duplicated and placed on the downstream end of the transposon, resulting in the following configuration: A-N-IR-*kanR*-IR-N-T. This detailed pattern indicates not only the sequence preference but also the consistency of the transposition event, ensuring the correct integration of the *kanR* cassette. Given, the successful transposition of these *IS5* constructs and their ability to mobilize cargo genes, we conclude that the active *cin* operon was laterally transferred via into *Orientia*.

## Discussion

*Wolbachia’s* CI operons (*cifs*) are mechanistically diverged into *cid*, *cin*, or *cnd* functional groups.^4^ CidB proteins induce CI via deubiquitylating activity and CinB proteins via nuclease activity.^1,5^ CndB proteins putatively induce RP via some mixture of both; though *cnd* operons are the least studied. Understanding how diverse *cifs* converged on or split into *cid*, *cin*, and *cnd* mechanisms will contribute to understanding how RP evolves and exhibits divergent phenotypes under different genetic contexts. Distant outgroups or edge cases can be most informative. To that effect, we sought to find and study a *cif* operon outside wolbachiae. Herein we demonstrated that an active *cinA/B* operon, capable of CI induction and rescue in *Drosophila melanogaster*, has jumped into the genome of a lethal human pathogen. The mechanism of lateral gene transfer can most parsimoniously be attributed to the immediately adjacent *IS*5 transposon, which bears IRs flanking the operon – constructing a larger mobile element. Herein we demonstrated the capability of this transposon to jump and introduce cargo, defining its mechanistic features. Thus, our data provides direct laboratory evidence demonstrating what was previously speculated based on genomic inferences^18,21^.

We hypothesize that jumping *cifs* facilitate the transfer of RP phenotypes to recipient bacteria. Downstream, this directly transfers gene-drive capabilities or at least promotes the evolution of gene-drive. Thus, our data suggest the obvious possibility that *O. tsutsugamushi* might be capable of pathogen gene-drive in mites. However, CI and/or RP due to this *cinA/B* operon hasn’t yet been fully characterized in *Leptotrombidium* mites, the vectors of scrub typhus.

We speculate that *cin/cnd* functions might explain multiple RP phenotypes including both CI and parthenogenesis (under certain mechanisms). Several lines of evidence support this. *Wolbachia* strains that infect the plant hopper *Laodelphax striatellus* induced strong CI ^77,78^. Genome sequencing revealed *w*Str has only a *cnd^w^*^Str^ operon ^21,58^. Given that *cid* and *cin* operons both induce CI,^1,5^ the best hypothesis is that *cnd^w^*^Str^ induces CI. Intriguingly *cnd* operons are also found on plasmids in parthenogenesis-inducing *R*. *felis* in booklice.^21^ It could be possible that *cnd* operons are capable of both CI and parthenogenesis under divergent contexts, or may be limited in scope when expressed in artificial systems. Similar thought applies directly to *cin*^*o*Tsu^. When expressed as a transgene in *D. melanogaster*, *cin*^*o*Tsu^ induces CI, yet the organism that it belongs to, *O. tsutsugamushi*, induces parthenogenesis in *Leptotrombidium* mites ^40,41^. Intriguingly, the CI data of this particular *cin* suggest that it has not evolved sex specific toxicity indicative of mature CI systems. The challenge will be studying these *cif* variants in their native systems and developing methods for determining if *cifs* are capable of inducing non-CI phenotypes.

Exploring if and how CI is connected mechanistically to other RP phenotypes like parthenogenesis will be a goal of future research. Thelytokous parthenogenesis is the development of females from unfertilized eggs. Several distinct mechanisms for this process have been described, including: (a) *Haploid female parthenogens*, where paternal chromosomes are destroyed or never merge with eggs, thereby producing haploid females^41^; (b) *Diploid female parthenogens*, where paternal chromosomes are destroyed or never merge, there is a signal to fuse (i.e., cellular/terminal fusion), and two post-meiotic eggs fuse to restore diploidy (*automictic thelytokous parthenogenesis* – mechanism 1 ^40^); (c) *automictic thelytokous parthenogenesis* – mechanism 2, wherein instead of fusion, a post-meiotic egg is replicated/duplicated to produce a diploid female parthenogen; and finally, (d) *apomictic thelytokous parthenogenesis*, where eggs that were never fertilized divide mitotically and diploid females produced are clones of their mother.

Thus the scientific term, parthenogenesis, includes a vast diversity of discrete molecular mechanisms that are all quite different – and there is even more complexity. With gynogenesis – a form of parthenogenesis in the viviparous poeciliid fish, paternal chromosomes are required for egg activation, but the paternal chromosome dissolves before it fuses with the egg nucleus.^79^. Even more bizarre, pseudogamy – a form of parthenogenesis in the beetle, *Ptinus clavipes* mobilis Panzer, paternal chromosomes are required to enter into the egg nucleus before eggs can develop parthenogenetically ^79^. Despite the sperm-dependencies of these additional examples of parthenogenesis, the paternal material never fuses with the egg.

One of these diverse mechanisms, *haploid female parthenogens*, might be explained by *cinA/B* operons. In a reported case of haploid female parthenogenesis in mites, mating occurs, a spermatophore is transferred, yet a symbiont destroys the paternal chromosomes. The embryo then resorts to default development as female. Destruction of paternal chromosomes would be within the capabilities of a *cinB*-like nuclease and therefore a CI inducer might also induce parthenogenesis in this context. All other forms of parthenogenesis, where male chromatin is targeted and destroyed, might be explained by a more specialized *cinB* nuclease. In fact, parthenogenesis itself might evolve from progressive tweaking of a guided nuclease to recognize specific epigenetic markers characteristic of males or females. It has been postulated that CI evolved from broad spectrum TA systems that select for plasmids.^4,21,80^ For this evolution to occur, the broad-spectrum toxin must become specific to male chromosomes alone. Permutating tandem nuclease domains (nuc_1_ and nuc_2_) to recognize or ignore epigenetic signals might be the very means to reach the terminal CI phenotype in arthropods. Thus, targeting mechanisms of *cinB* nucleases and specific differences encoded in the tandem nuclease domains remains an important focus of future studies.

Overall, our work here demonstrates CI capabilities in an operon outside of wolbachiae and evidences a novel mode of lateral gene transfer of that very operon. Further, our study bolsters prior reports that implemented bioinformatics and evolutionary analyses to identify a broad range of *cif* operons in many divergent intracellular bacteria, as well as some of the host genomes they parasitize ^18,21^. To our knowledge, this characterization of a non-*Wolbachia cif* operon from another obligate intracellular bacterium vectored by arthropods is the first demonstration that “*a veritable cornucopia of architecturally diverse proteins found almost exclusively in a narrow range of intracellular bacteria*” is capable of exerting diverse RP phenotypes ^21^.

This has profound impact on vector biology, as the very initiatives to utilize wolbachiae-infected mosquitoes to outcompete mosquitoes infected with deadly human viruses might apply to other systems where arthropods transmit blood-borne pathogens to humans ^81,82^. Thus, the utility of the *Wolbachia cif* gene drive mechanism to combat vector-borne human microbial pathogens can be expanded to a wide range of blood-feeding arthropods that often harbor a bevy of endosymbionts – by simple transfer of *cifs*. Additional bioengineering these endosymbionts with genes encoding factors that destroy incidental human pathogens is a path toward eradicating vector-borne human diseases, which remain a significant cause of human death globally ^83^.

## Materials and Methods

### *O. tsutsugamishi* Genome Analysis (Joes Part)

A recent study devoted great effort to the identification of RAGE and inter-RAGE regions in the genomes of eight *O. tsutsugamishi* strains (Ikeda, Boryong, Karp, Kato, Gilliam, TA686, UT76 and UT176)^60^. Criteria for sequence analysis, detailed in that study, led to identifying and numbering inter-RAGE regions within these genomes. RAGE regions were subsequently characterized using information largely drawn from Nakayama *et al*. ^51^, with minor modification to annotations. The Ot *cinA*/*B* operon, identified previously ^21^, was used in blastp searches against the eight Ot strains to establish the precise genomic positions relative to RAGE and inter-RAGE regions. All CinA and CinB proteins from each *O. tsutsugamishi* genome were inspected for features defining CinA and CinB proteins from other organisms, particularly the dual nuclease active sites.

To determine the similarity of the Ot CinA and CinB proteins to analogs from other organisms, we used blastp to query the NCBI nr database for the best matches to each sequence (e-value < 1, BLOSUM45 matrix). Hits were downloaded and then binned by taxonomic assignment as either bacterial (“Rickettsia”, “Wolbachia”, “Cardinium”, or “other”) or eumetazoan (“Rotifer”, “Coleoptera”, “Araneae”, “Hemiptera”, and “Hymenoptera”). All hits were then subsequently plotted by descending Bit score. For protein models, %ID was determined by blastp alignments, as well as more refined alignments using MUSCLE v3.8.31 (default parameters) ^84^.

### *Drosophila* Lines and Maintenance

Transgenic flies were generated with ΦC31 site-specific *attP*/*B* site integrations onto chromosome 3 of fly (BDSC #9744). Microinjections were contracted to BestGene Inc. Integrations were confirmed by multiple methods including a mini-white cassette (red eye marker genes), CI induction/rescue phenotypes, and in some cases via PCR amplification of the *attL* recombination site (performed by BestGene Inc). Fly line MTD-Gal4 (gift from Lynn Cooley, Yale University) was used as the driver for all UAS-transgenes. Fly lines were verified to be free of *Wolbachia* infection via PCR amplification of the *virD4* gene with *Wolbachia*-specific primers. Fly lines were maintained on cornmeal-glucose-based solid media at 23°C with an ∼12-hour light-dark cycle.

### CI Construct Assembly and Manipulation

*cinA^oTsu^* and *cinB^oTsu^* were initially cloned from genomic *Orientia tsutsugamushi* (Ikeda strain) DNA into a gateway vector pBluescript SK (+) (pBSK) using PCR and restriction enzyme cloning (see Construct Database). PCR primers are listed in Supplemental Table 1. In general, PCR used Phusion HF DNA polymerase (New England Biolabs). Amplicons were purified by E.Z.N.A.® Cycle-Pure Kit (Omega Bio-Tek), restriction digested with various enzymes (New England Biolabs), agarose gel-purified with E.Z.N.A.® Gel Extraction Kit (Omega Bio-Tek), and ligated into pBSK. Nuclease permutation mutants were generated by site-directed mutagenesis using pBSK_*cinB* as a template. These mutants include pBSK_*cinB*^oTsu^(Off – Off), pBSK_*cinB*^oTsu^(On – On), and pBSK_*cinB*^oTsu^(Off – On). Operon constructs bearing the T2A peptide insertion were codon optimized and synthesized by Genscript. These constructs include pUC57_*cinA/B*^oTsu^(On – Off) and pUC57_*cinA/B*^oTsu^(Off – Off). All genes were subcloned into pUASp-*attb* with 5′ NotI-HF and 3′ BamHI-HF. Due to the size similarity of *cinB* and the gateway backbone, subcloning sometimes also included an additional enzyme BglI to remove the gateway backbone.

### Hatch Rate Analysis

F₀ crosses with male UAS-transgene and female MTD-Gal4 were set up in fresh food vials, and later adults were removed. From these progeny, virgin F₁ males (MTD-Gal4;UAS-transgene) were collected during a 2-day period and aged to 3–4 days. When collecting virgins, flies were kept at 18°C overnight and at room temperature during the day, with periodic collection periods not greater than 16 or 8 hours, respectively. Individual virgin F₁ aged males (MTD-Gal4;UAS-transgene flies) were pairwise crossed to virgin uninfected (U; wCS) flies on apple juice plates smeared with a dab of yeast paste. Flies were allowed to mate and lay eggs for 36 hours, then apple juice plates were changed. Flies were allowed to lay eggs for an additional 24 hours. Afterward, the adults were removed, and plates were incubated for 36 hours to allow eggs to hatch. Hatched and unhatched eggs were counted to calculate hatch rate. Plates with fewer than 10 eggs were excluded from statistical analyses for hatch rate. Fecundity was calculated from the sum of hatched and unhatched eggs without removing plates with fewer than 10 eggs.

### RNA Isolation and RT-qPCR

Total RNA was extracted from *Drosophila* eggs (200-300 eggs, 12-24 hours after egg deposition) using the E.Z.N.A. Total RNA Kit I (Omega Bio-Tek, R6834-01), following the manufacturer’s protocol. Eggs were placed in bead-ruptor tubes containing acid-washed beads and 350 µl of lysis buffer. Homogenization was carried out using the Bead Mill Homogenizer (OMNI International), and the supernatant was transferred into clean 1.5 ml Eppendorf tubes. Ethanol (70%) was added at a 1:1 ratio before transferring the sample to HiBind RNA Mini columns in 2 ml collection tubes. After centrifugation (1 min) the columns were subjected to DNase I digestion to remove genomic DNA. Columns were then washed with RNA Wash Buffer I, then II, then centrifuged (10,000 x g). RNA was eluted in 40 µl of nuclease-free water and stored at −80 °C. The purity of RNA was assessed using spectrophotometry (A260/A280 ratio). cDNA synthesis was performed using 1 µg of total RNA with the SuperScript III Reverse Transcriptase enzyme (Thermo Fisher Scientific) in a 20 µl reaction volume, according to the manufacturer’s instructions. Quantitative real-time PCR (RT-qPCR) was conducted using the Bio-Rad CFX96 Real-Time PCR system, with SsoAdvanced Universal SYBR Green Supermix (Bio-Rad) in a 10 µl reaction volume. Gene expression was measured in technical duplicates, and mRNA levels were normalized to the *Actin 5C* (CG4027) housekeeping gene^85^. Reactions were performed in 96-well plates primers listed in Supplemental Table 1. Changes in gene expression levels were calculated using the 2^-ΔΔC^ method ^86^.

### Microscopy and Supplemental Images

2–4 day old F₁ females (MTD-Gal4;UAS-transgene) were placed on a microscope slide with 1X PBS. Ovaries were dissected and identified by tracing the ovarioles to the lateral and common oviduct. Ovaries were transferred to a clean microscope slide and imaged on a brightfield dissecting scope (Nikon SMZ1270). Afterward, ovaries were stained with DAPI (Biotium) (10 µg/mL in 1X PBS) and incubated at room temperature for 5 min, then imaged on a fluorescent microscope (Nikon Ts2R). Images were contrast-adjusted in Adobe Photoshop.

### *E. coli* Strains and Culture Conditions

*Escherichia coli* strains used in this study included *Top 10 F’* and *BL21 AI. Top 10 F’* strains were utilized as storage hosts for developed constructs, while BL21 AI strains were used as the heterologous protein expression strain compatible with arabinose induction used in the pBAD system. Chemically competent cells were prepared and transformed using standard chemical transformation methods as described by ^87^. Unless otherwise stated, *E. coli* cultures were grown in Luria Broth (LB) supplemented with Ampicillin (AMP) (50 µg/mL) at 37°C.

### Counter-Selective M63 Media

For DOG selection experiments, a counter-selective M63 minimal medium was prepared with a standard recipe. To prepare the 5X M63 stock solution, the following reagents were added to 750 mL of sterile distilled water in a 2 L flask: 10 g ammonium sulfate ((NH₄)₂SO₄), 68 g potassium dihydrogen phosphate (KH₂PO₄), and 2.5 mg ferrous sulfate heptahydrate (FeSO₄·7H₂O). The components were dissolved with heating and stirring, followed by pH adjustment to 7.0 using phosphoric acid (H₃PO₄) (to increase pH) or potassium hydroxide (KOH) (to decrease pH). The final volume was adjusted to 1L with sterile distilled water, and the solution was autoclaved. The counter-selection process using DOG followed the method outlined by Barken et. al. ^88^. For the preparation of counter-selective M63 agar plates, 200 mL of the 5X M63 stock was diluted with 789 mL of sterile distilled water and combined with 20 g agar. The mixture was autoclaved and cooled to 50°C before adding the following sterile-filtered supplements: 1 mL of 1M magnesium sulfate heptahydrate (MgSO₄·7H₂O), 2.668 mL of 75% glycerol (final concentration: 0.2%), 0.1 mL of 0.5% thiamine, 1 mL of Kanamycin (50 mg/mL), and 10 mL of 20% DOG (final concentration: 0.2%). Plates were poured and allowed to dry for 48 hours before use.

### Transposon Gene Classifications, Sources, and Constructs

The *IS5* transposon constructs in this study contained three key cloned genes, which are the *IS5* transposon, the *galK* counterselection cassette, and the *kanR* cargo gene flanked by inverted repeats (**Sup Fig 4**.). There were multiple homologous transposon constructs we analyzed in this study. The first was a wild type sequence of *IS5* from *O. tsutsugamushi* str. IKEDA with minor modifications. Specifically, the native WT gene in the *Orientia* genome is pseudogenized and likely inactive due to three premature stop codons. We cloned the WT sequence from genomic *Orientia* DNA into pBSK and repaired the three premature stop codons using site-directed mutagenesis via the QuikChange protocol. Secondly, A synthetic variant with the same repaired amino acid sequence was synthesized and codon optimized by Genscript. These two transposon configurations were experimentally compared in **Figure 5a**. The *galK* and *kanR* genes were cloned from pgalK and pET-28pp plasmids, respectively.

**Figure 5.**
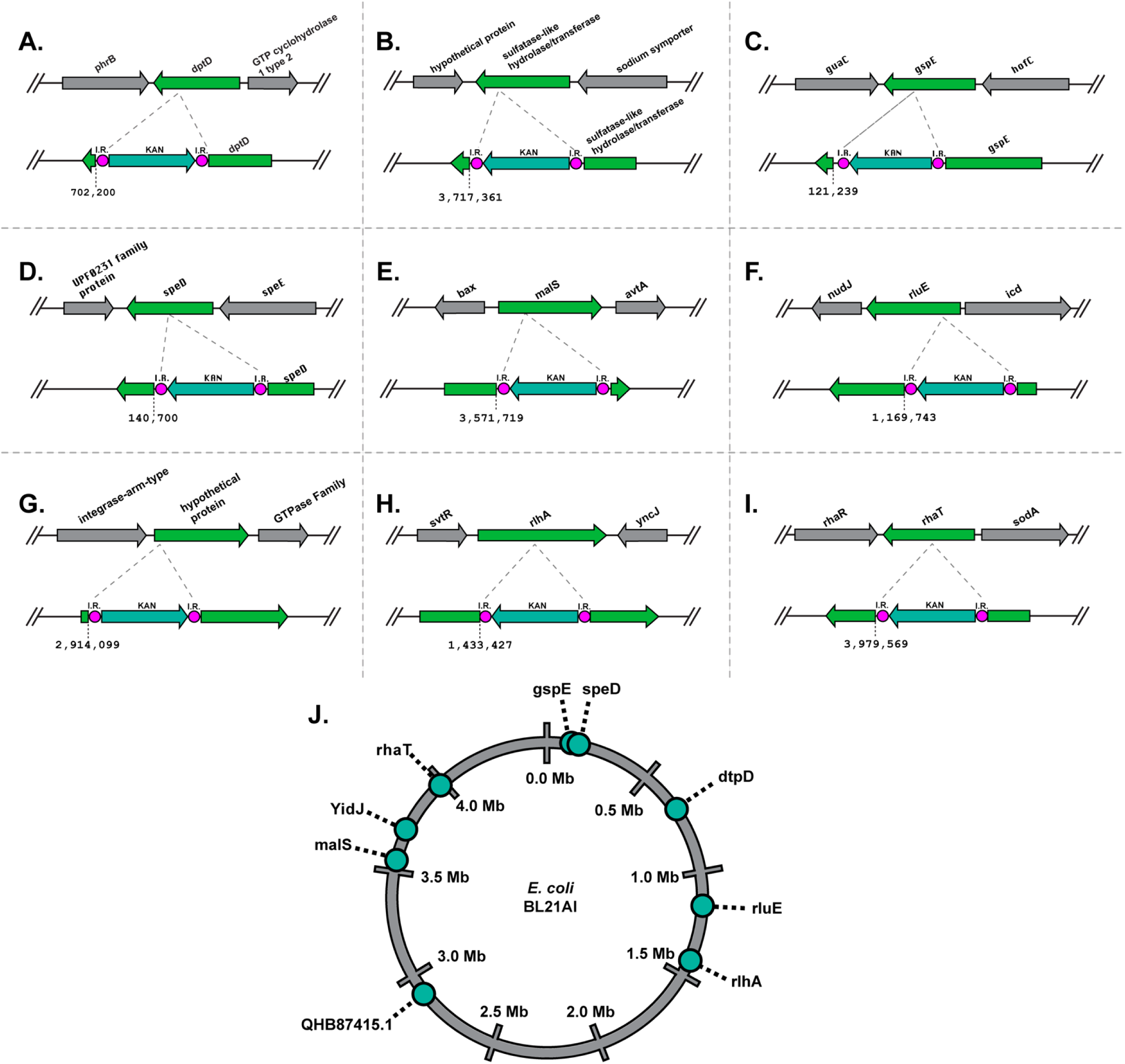
*kanR* cassette integration sites in the *E. coli* BL21-AI genome. The figure shows ten loci (**a**–**j**) where the *kanR* cassette was integrated via transposon activity, mapped against the *E. coli* BL21-AI reference genome (NCBI accession number CP047231.1). Integration occurred within **a.** the dipeptide permease D (*dtpD*) gene at position 702,200; **b.** the sulfatase-like hydrolase YidJ gene at 3,717,361. **c.** the ATPase component of the type II secretion system gene (*gspE*) at 121,239; **d.** the S-adenosylmethionine decarboxylase proenzyme (*speD*) gene at 140,700; **e.** the α-amylase (*malS*) gene at 3,571,719; **f.** the 23S rRNA pseudouridine^2457^ synthase (*rluE*) gene at 1,169,743; **g.** a hypothetical protein encoding gene (QHB87415.1) at 2,914,099; **h.** the 23S rRNA 5-hydroxycytidine C2501 synthase (*rlhA*) gene at 1,433,427; and **i.** the rhamnose/lyxose:H^+^ symporter (*rhaT*) gene at 3,979,569. Dashed lines mark insertion points; green arrows indicate the *kanR* cassette; grey arrows represent flanking genes; pink circles highlight inverted repeats. **j.** Circular genome map of inserts on E.coli BL21AI.

To examine the influence of inverted repeat (IR) length on transposition efficiency, we constructed two versions of the *kanR* cassette flanked by IRs of different lengths. The long IRs (117 bp) correspond to the native IR sequences from the *O. tsutsugamushi IS*5 transposon and were synthesized along with the *kanR* cassette by Genscript. The short IRs (17 bp) were designed based on conserved core sequences within the native IRs and were incorporated directly into the PCR primers used to amplify the *kanR* cassette, facilitating seamless cloning. This approach allowed us to assess whether the full-length native IRs were necessary for transposase recognition and activity or if shorter IRs could effectively mediate transposition.

With respect to plasmid assembly, the order of gene insertion varied depending on construct being built, but the insertion sites remained consistent. Transposons without IRs were inserted downstream of the arabinose-induced promoter, between 5’ NcoI and 3’ XhoI sites. The *galK* gene, consisting of its open reading frame (ORF) and constitutive promoter, was inserted at the 5’ XhoI and 3’ PstI sites. The *kanR* cassette, also with its constitutive promoter, was inserted at the 5’ PstI and 3’ HindIII sites, with terminal IRs and a small multi-cloning site (5’ KpnI and 3’ EcoRI) added between the 3’ end of *kanR* and the 5’ end of the right IR. Short inverted repeats were simply added into the PCR primers (see Supplemental Table. 1). In contrast, long IRs were synthesized conjoined to a *kanR* cassette by Genscript. All primers and genes used in construct design are Supplemental Table1. and enzymes were purchased from New England Biolabs.

### PCR, Cloning Procedures, and Plasmid Validation for Transposon Constructs

PCR cloning protocols followed as described above in CI construct assembly and manipulation. Once purified, the PCR products were ligated into plasmid vectors, such as pBAD (ThermoFisher) for arabinose induction and pBluescript SK+ (Agilent Technologies, USA). Plasmid inserts were validated by restriction enzyme digest analysis and Sanger sequencing. Sanger sequencing was performed by MC Labs using ABI 3730XL sequencers, while whole plasmid sequencing was conducted by Plasmidsaurus, utilizing Oxford Nanopore Technology with custom analysis and annotation. Point mutations were introduced using standard PCR based site directed mutagenesis protocols, and larger modifications were carried out using the site-directed, ligase-independent mutagenesis (SLIM) method ^89^.

### Experimental Setup and Successful Counterselection of galK-Positive Plasmids with 2-deoxy-D-galactose

To assess the jumping capabilities of the transposons and evaluate the efficacy of DOG in our constructs, BL21 AI strains were transformed with the transposon constructs containing a 1,179 bp segment of the *E. coli*-derived *galK* gene. The BL21 AI strain lacks the *galK* gene, allowing us to use it as a counterselection marker within experiments.To start overnight (O/N) cultures were inoculated in LB with Ampicillin and incubated at 37°C. The following day, the O/N cultures were diluted into 200 mL cultures and grown to an OD₆₀₀ of 0.5, measured using a spectrophotometer (UV-1600PC). Cultures were then split into two 100 mL aliquots: one for the positive control (POS), to which arabinose was added to a final concentration of 0.2%, and one for the negative control (NEG) without arabinose. The cultures were incubated at 37°C for approximately 6 hours in a shaking incubator set to 200 rpm (Thermofisher Scientific™, MaxQ™4000) to induce transposase expression, facilitating the mobilization and integration of the *kanR* cassette into the genome. After incubation, cells were pelleted by centrifugation at 5,000 rpm for 10 min, and the supernatant aspirated. The pelleted cells were then resuspended and washed twice with sterile distilled (DI) water, taking care to minimize cell loss due to poor adhesion. The cell suspensions were standardized to a concentration of approximately 10⁸ cells/mL (∼1.0 OD), and 100 µL (∼10⁷ cells/plate) were plated in triplicate on M63 counter-selective media containing 0.2% DOG, 0.2% glycerol, and Kanamycin. The *galK*-positive cells convert DOG into 2-deoxy-galactose-1-phosphate, a toxic intermediate that impairs cell viability, enabling successful counterselection against *galK*-positive plasmids (Fig. 5b). Plates were incubated at 37°C for 36 hours. Colony-forming units (CFUs) were counted on both positive and negative plates, and the results were graphed and analyzed for statistical significance. This method enabled effective plasmid expression and facilitated the selection of *kanR* insertions into the host genome, confirming the utility of DOG as a counterselective agent. This trial was done in triplicate.

### Transposon Validation Procedure

If CFU counts in the positive arabinose sample were significantly higher than in the negative control (p < 0.05) indicating transposon mobilization, further analyses were conducted to verify transposon insertions. When the positive samples displayed significance, five colonies were randomly selected from each of the three plates (15 colonies total for each condition, positive and negative). These colonies were cultured overnight in LB + Kanamycin (KAN) to maintain the cassette insertion. The following morning, plasmid DNA was extracted using the E.Z.N.A. Plasmid DNA Mini Kit I (Omega Biotech, D6942). Samples were analyzed on 0.8% agarose gels with ethidium bromide. Clones that had lost the plasmids from counter-selection were then sent to SeqCenter for genome sequencing. Sequencing was performed using the Illumina Whole Genome Sequencing pipeline, with DNA extracted using ZymoBIOMICS DNA extraction kits. The Illumina DNA Prep kit was used for library preparation, producing ∼280 bp inserts suitable for 150 bp paired-end read output. Libraries were indexed with IDT 10 bp UDI indices and demultiplexed bioinformatically after sequencing.

### Post-Sequencing Analysis

Contigs were assembled using BV-BRC’s bacterial genome assembly software (www.bv-brc.org/app/Assembly2) with Unicycler v0.4.8, utilizing default settings. The assembly was performed on normalized short-read data processed by Trim Galore (version 0.6.5dev) and BBNorm (October 19, 2017). Following assembly, two rounds of polishing were conducted using Pilon (version 1.23), and contigs were filtered based on length (minimum 300 bp) and coverage (minimum 5x) using Samtools (v1.17) and evaluated with Quast (v5.2.0). The resulting contigs were analyzed using BLAST to identify insertion regions, focusing on localizing the *kanR* cassette and its associated inverted repeats (IRs). These matches were cross-referenced against the BL21-AI annotated reference genome (CP047231.1) to determine the insertion sites and their locations.

### Statistical Analysis

For CI cross data, one-way ANOVA with multiple comparison was performed using Graphpad Prism. For RT-qPCR analyses, relative gene expression was quantified using the ΔΔCt method after normalization to an appropriate reference gene. Data normality for each target was first assessed, and as none of the datasets conformed to a normal distribution, a non-parametric approach was required. A Kruskal–Wallis test was used to evaluate overall differences among the experimental groups, followed by Dunn’s multiple comparisons test for pairwise analyses. A p-value of less than 0.05 was set as the threshold for statistical significance, and all analyses were performed using GraphPad Prism. For CFU analyses, A two-way analysis of variance (ANOVA) was conducted using GraphPad Prism (most recent version) to examine the effects and interactions of two independent variables on the dependent variable. This method allowed for the assessment of each variable’s significance and strength, as well as any interaction effects. A P-value of less than 0.05 was considered statistically significant.

## Acknowledgements

This work was supported by Auburn University startup funds for JFB, an National Science Foundation - CAREER AWARD (2335888) to JFB and United States Department of Agriculture - AFRI (2023-67016-39659) to JFB. Throughout the duration of this project, JJG was supported by funds from the National Institute of Health/National Institute of Allergy and Infectious Diseases grants R21AI26108 and R21AI166832.

## Statements about data availability

All data are available in the manuscript and supplemental materials.

## Author Contribution

**JFB:** Supervision, Acquired funding, Designed experiments, performed experiments, wrote the manuscript, edited the manuscript. JJG: Supervision, Acquired Funding, Designed Experiments, Performed Experiments, wrote the manuscript, edited the manuscript. KO: Designed experiments, performed experiments, wrote the manuscript, edited the manuscript. SO: Designed experiments, performed experiments, wrote the manuscript, edited the manuscript. LM: performed experiments.

---

**Supplementary Figure 1.**
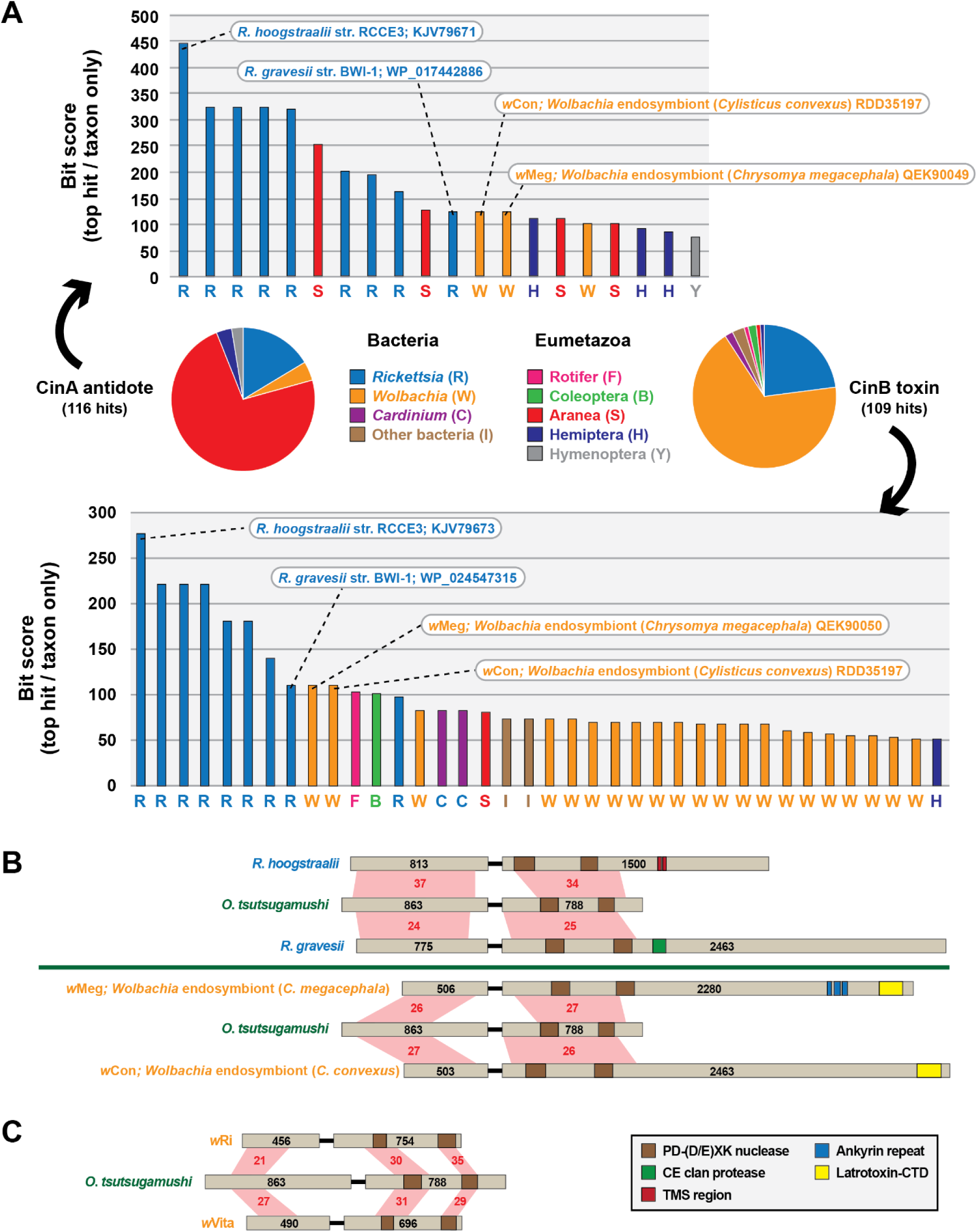
Phylogenomics analysis of *Orientia tsutsugamushi cin* operons **a.** Blastp results using CinA (top) and CinB (bottom) of *O*. *tsutsugamushi* str. Boryong as a query in searches against the NCBI nr protein database. Subjects are binned by taxonomy with the best hit per taxon shown in the graph. Operons shown in panel **b** are denoted. **b.** *Orientia cin* operons are similar to *Rickettsia* (top) and *Wolbachia* (bottom) *cin* or *cnd* operons that carry large multi-domain toxins. **c.** *Orientia cin* operons are more divergent than the described CI-inducing *cin* operons of wolbachiae. Black numbers on proteins depict sizes (aa). Red shading shows %ID (aa).

**Supplemental Figure 2.**
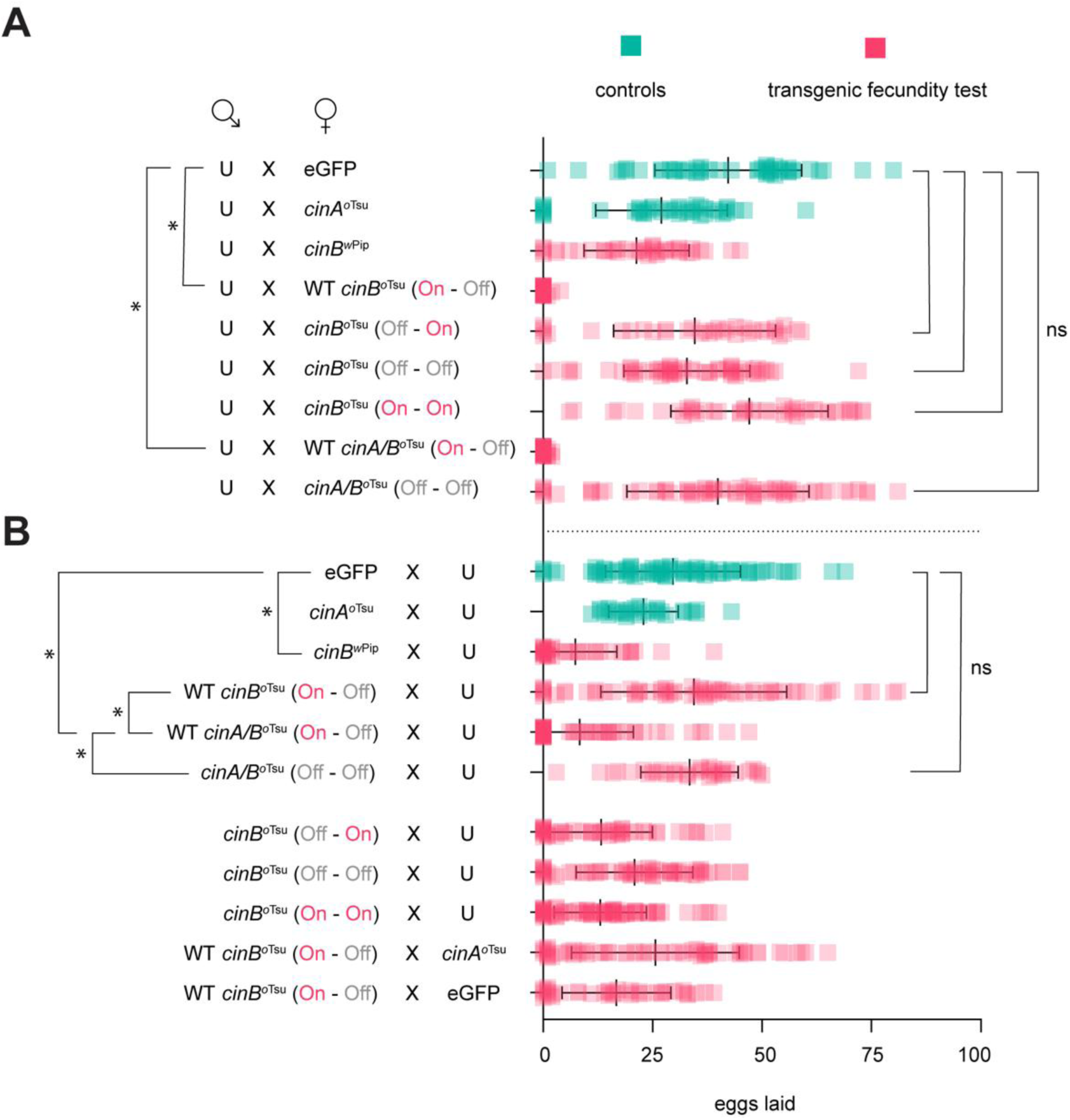
Fecundity analysis of transgenic *cin*^*o*Tsu^ constructs. Green data are control experiments and pink data are novel transgenic construct tests. **a.** Crosses of uninfected (U) males mated to females expressing transgenes. **b.** Crosses with transgenes in males mated to uninfected mothers; or mothers with *cin*A^*o*Tsu^ or eGFP in rescue experiments. Lines are means with standard deviation. Significance (*) is p < 0.0001 and ns is not significant by ANOVA with multiple comparison between all groups and Tukey’s post-hoc analysis. GAL4/UAS driver is MTD.

**Supplemental Figure 3.**
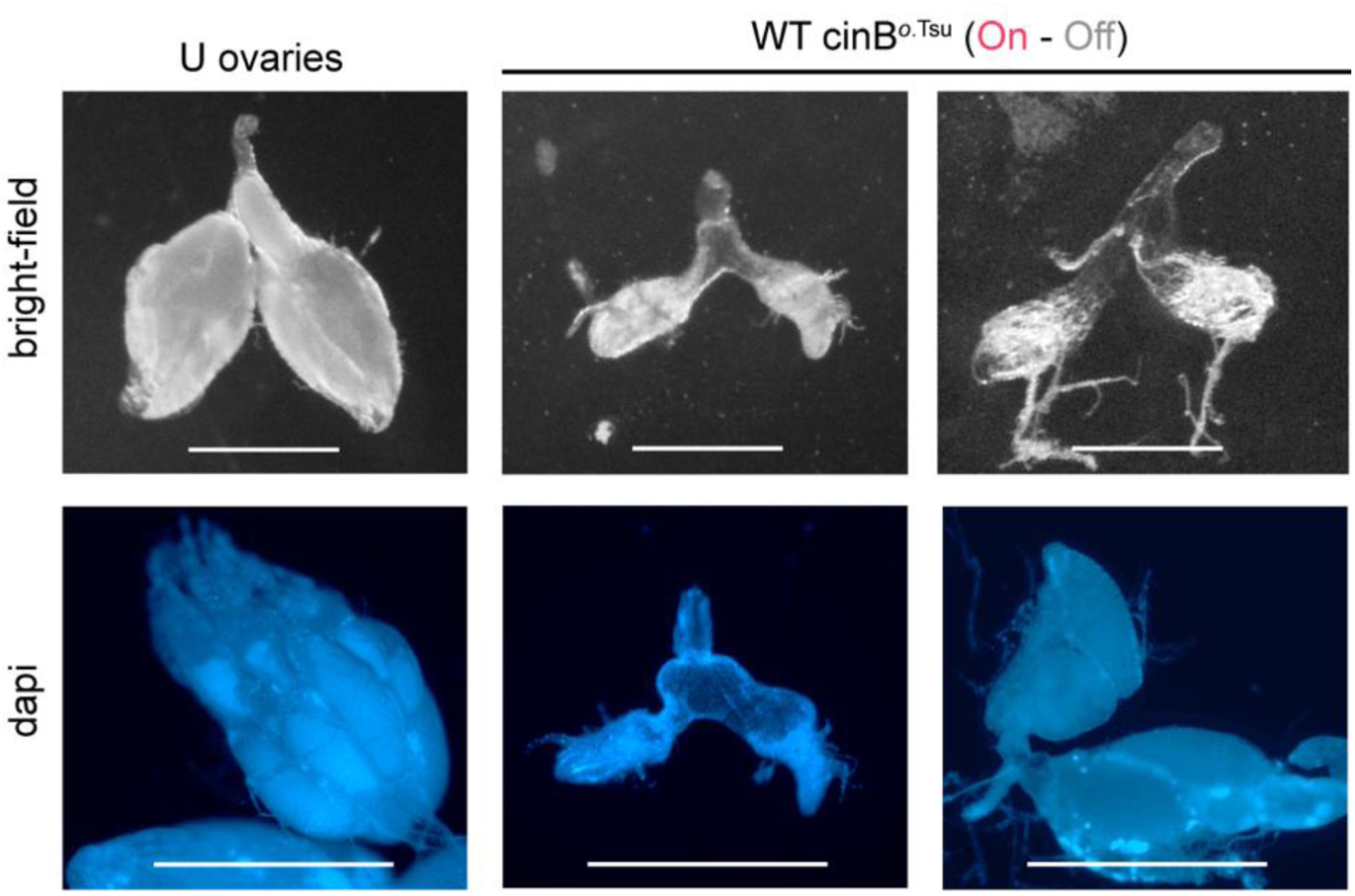
Microscopy of fruit fly ovaries. At left are normal uninfected ovaries (U). At right are ovaries expressing a WT *cin*B^oTsu^ (On - Off) transgene with the MTD driver. Ovaries show gross morphological defects including stunting, abnormal oviducts, and asymmetry. Scale bars are 0.5 mm. Ovaries are representative of 60 ovaries viewed from transgenic females.

**Supplemental Figure 4.**
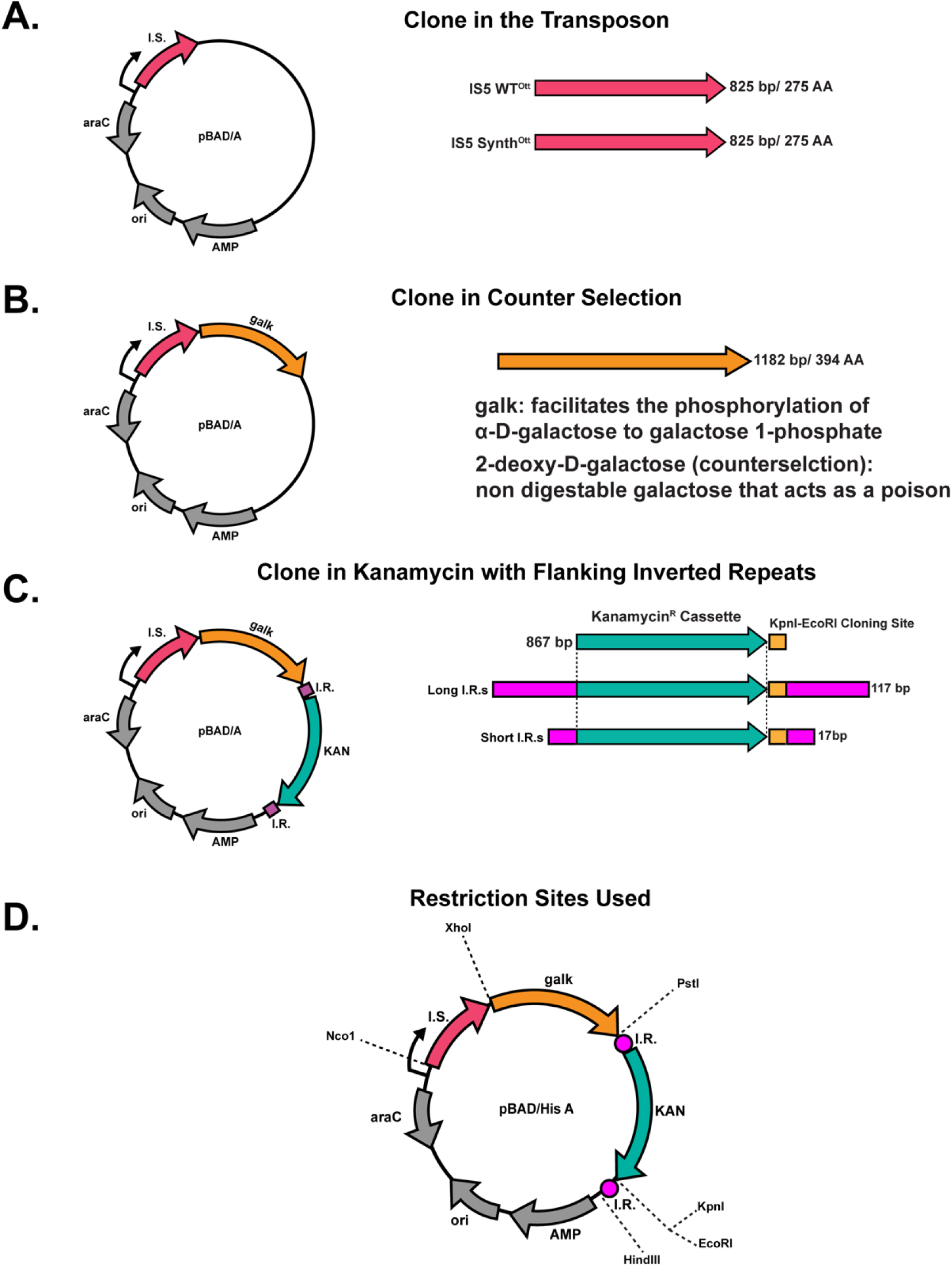
Stepwise construction of plasmid constructs using the pBAD/A backbone. Each panel (a–d) represents a stage in the construct design process, detailing gene insert locations, variants used and insert sizes. **a.** Integration of transposon elements into the pBAD/A backbone; the two transposon genes encode identical amino acid sequences. **b.** Incorporation of the *galK* gene (1,182 bp, encoding 394 amino acids) as a counter-selective marker, facilitating the phosphorylation of α-D-galactose to galactose-1-phosphate. **c.** Addition of the *kanR* gene, flanked by two distinct inverted repeat pairs: Long IRs derived from wild-type sequences and Short IRs synthetically derived from the wild-type sequence. A downstream cloning site enables the insertion of extra genetic material. **d.** Final schematic of the complete construct with annotated restriction sites used in each step.

**Supplementary Figure 5.**
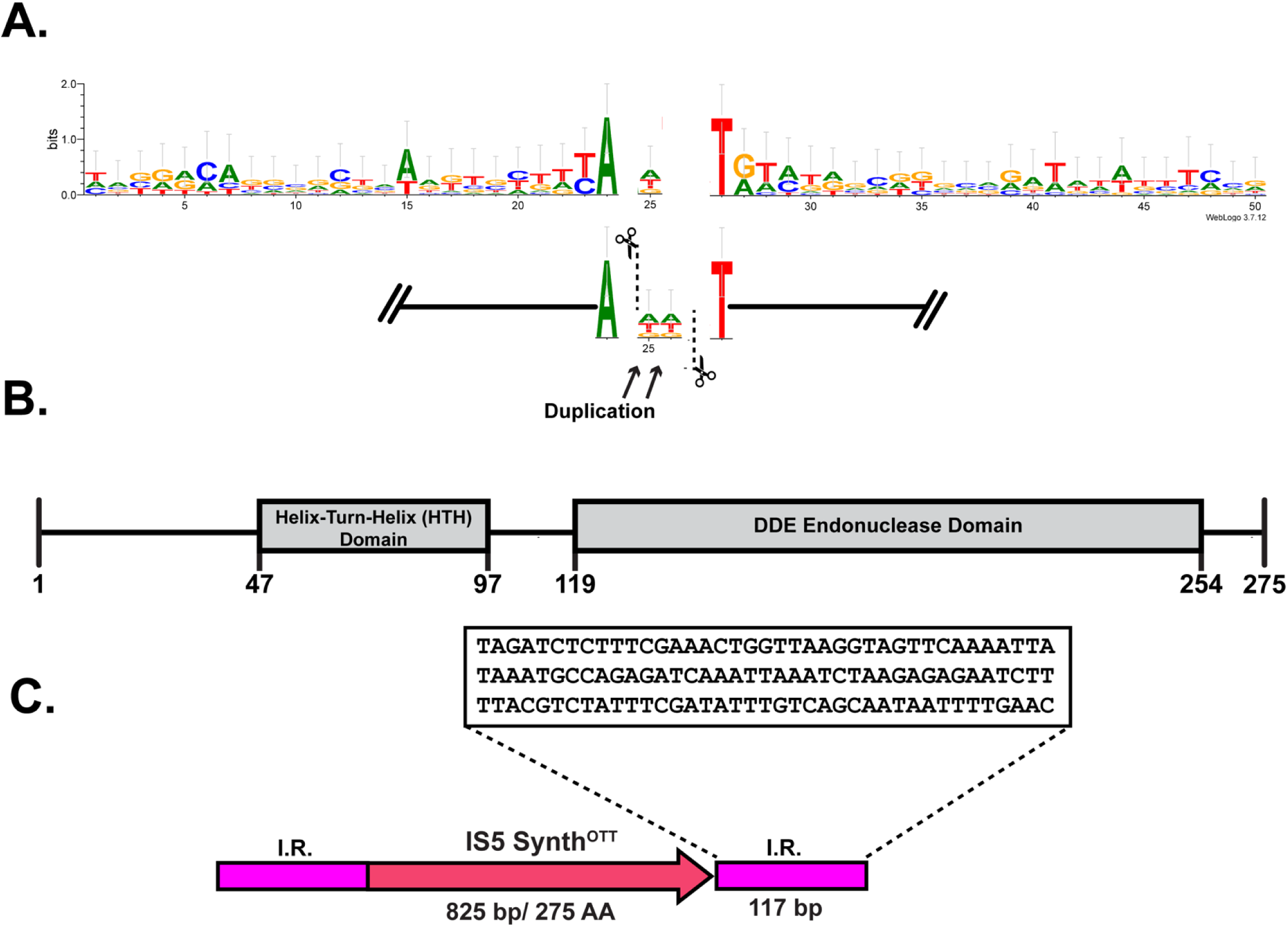
Functional analysis of transposon insertion and associated sequence elements in *E. coli* BL21-AI**. a.** Sequence logo of nucleotide composition and conservation around the insertion sites, generated using WebLogo 3.7.12 from sequences flanking 10 transposon insertion events. Position 25 marks where direct repeats form post-insertion. The lower schematic illustrates the insertion event, with arrows indicating DNA cleavage points and the direct repeats formed at the insertion site. **b.** Domain architecture of the synthetic *IS*5 SynthOTT transposase, predicted using InterPro. **c.** Structure of the *IS*5 SynthOTT transposon with inverted repeats.

**Supplementary Figure 6.**
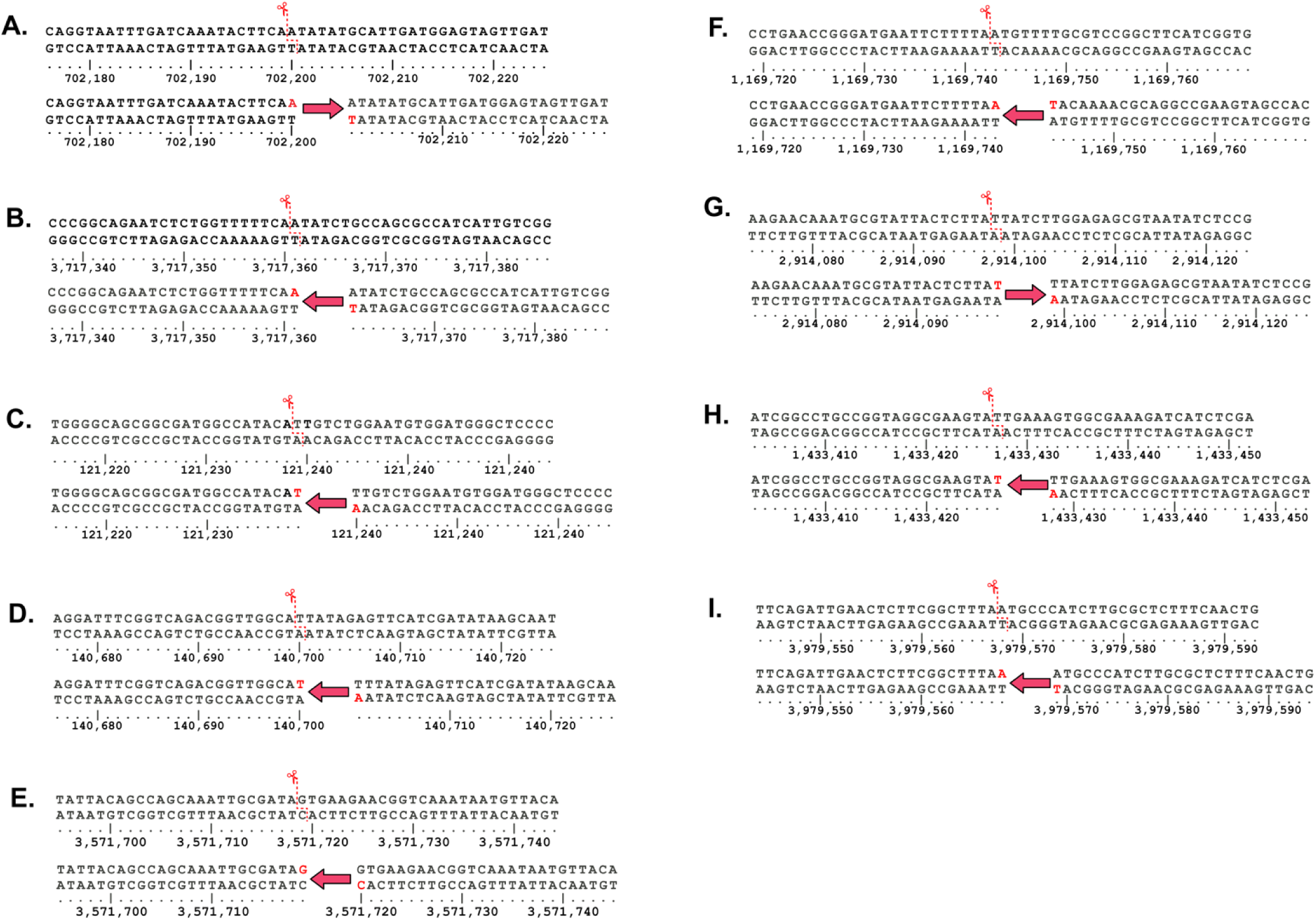
Exact sequence flanking *kanR* cassette integration sites in the *E. coli* BL21-AI genome. The figure displays the 25 bp sequences upstream and downstream of ten loci (a–j) where the *kanR* cassette was integrated via transposon activity, driven by the *IS*5 Synth OTT transposon protein, mapped against the *E. coli* BL21-AI reference genome (CP047231.1). Integration occurred within (a) the *dtpD* gene at position 702,200, (b) the *yidJ* gene at position 3,717,361, (c) the *gspE* gene at position 121,239, (d) the *speD* gene at position 140,700, and (e) the *malS* gene at position 3,571,719. Additional insertions occurred in (f) the *rluE* gene at position 1,169,743, (g) a hypothetical protein encoding gene (QHB87415.1) at position 2,914,099, (h) the *rlhA* gene at position 1,433,427, and (i) the *rhaT* gene at position 3,979,569. Red arrows indicate the direction of insertion, and red nucleotides show sequence duplications generated during transposition. The sequences are annotated with genomic coordinates, with numbers underneath indicating their positions relative to the *E. coli* BL21-AI genome.

**Supplementary Figure 7.**
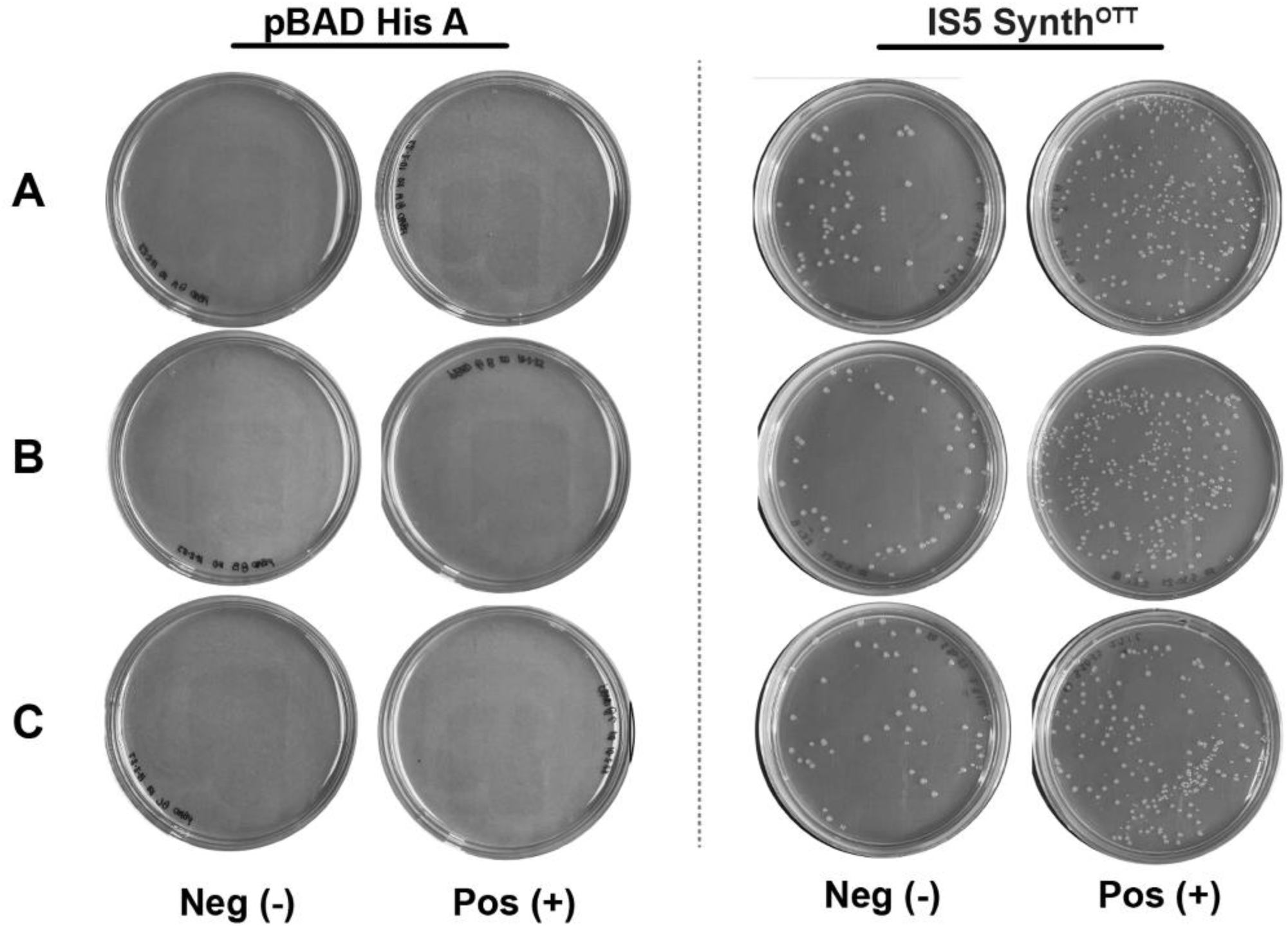
DOG trial comparing transposon activity in *E. coli* BL21-AI. Growth of *E. coli* BL21-AI on M63 media with Kanamycin. The left two columns show pBAD His A (blank control), and the right two columns show *IS*5 SynthOTT. Plates were incubated under non-induced (Neg -) and arabinose-induced (Pos +) conditions. Rows A-C represent biological replicates from a single trial.

